# Tenascin C Deletion Impairs Tendon Healing and Functional Recovery After Rotator Cuff Repair

**DOI:** 10.1101/2024.09.11.612543

**Authors:** Robert Z. Tashjian, Jared Zitnay, Nikolas H. Kazmers, Shivakumar R. Veerabhadraiah, Antonio C. Zelada, Matthew Honeggar, Matthew C. Smith, Peter N. Chalmers, Heath B. Henninger, Michael J. Jurynec

**Author notes:** Corresponding Author: Robert Z. Tashjian, MD, University of Utah Orthopaedic Center 590 Wakara Way, Salt Lake City, UT 84108. All work was performed at the Department of Orthopaedics, University of Utah School of Medicine, Salt Lake City. **Disclaimer:** Robert Z. Tashjian is a paid consultant for Zimmer/Biomet, Stryker, Enovis and Mitek; has stock in Conextions and Genesis; receives intellectual property royalties from Stryker, Shoulder Innovations and Zimmer/Biomet; receives publishing royalties from the Journal of Bone and Joint Surgery and Springer and serves on the editorial board for the Journal of Bone and Joint Surgery. Peter Chalmers is a paid consultant for Depuy, Exactech, Smith and Nephew, and Enovis, serves on the editorial board for the Journal of Shoulder and Elbow Surgery, receives intellectual property royalties from Depuy, Exactech, and Responsive, and has equity in TitinKM. Jared Zitnay, Nikolas Kazmers, Shivakumar R. Veerabhadraiah, Antonio C. Zelada, Matthew Honeggar, Matthew Connor Smith, Heath Henninger and Michael Jurynec, their immediate family and any research foundation with which they are affiliated did not receive any financial payments or other benefits from any commercial entity related to the subject of this article. **IRB Approval: IRB Approval:** Animal work was performed in accordance with laws for the care and use of laboratory animals with approval by the University of Utah’s Institutional Animal Care and Use Committee (IACUC Protocol number 21-07004). **Author Contribution Statement:** Robert Tashjian and Michael Jurynec contributed substantially to the research design, data acquisition, data analysis, data interpretation, manuscript preparation, manuscript revision. Jared Zitnay, Heath Henninger and Nikolas Kazmers contributed significantly to the research design, data acquisition, data analysis, manuscript preparation and revision. Shivakumar R. Veerabhadraiah, PhD, Antonio C. Zelada, BS, Matthew Honeggar, BS, Matthew Connor Smith contributed significantly to the data acquisition and data analysis. Peter Chalmers contributed significantly to the research design, data analysis, manuscript preparation and revision.

## Abstract

The biological factors that affect healing after rotator cuff repair (RCR) are not well understood. Genetic variants in the extracellular matrix protein Tenascin C (*TNC*) are associated with impaired tendon healing and it is expressed in rotator cuff tendon tissue after injury, suggesting it may have a role in the repair process. The purpose of the current study was to determine the role of TNC on tendon healing after RCR in a murine model. The supraspinatus tendon was transected and repaired on the left shoulder of Wild-Type (WT-RCR), *Tenascin C* null (*Tnc^−^-*RCR) and *Tnc* heterozygous (*Tnc*^+/−^-RCR) mice. Controls included the unoperated, contralateral shoulder of WT-RCR, *Tnc^—^*RCR, *Tnc*^+/−^-RCR mice and unoperated shoulders from age and genotype matched controls. We performed histologic, activity testing, RNA-seq, and biomechanical analyses. At 8-weeks post-RCR, *Tnc^−^* and *Tnc*^+/−^ mice had severe bone and tendon defects following rotator cuff repair. *Tnc^−^*-RCR mice had reduced activity after rotator cuff repair including reduced wheel rotations, wheel duration, and wheel episode average velocity compared with WT-RCR. Loss of *Tnc* following RCR altered gene expression in the shoulder, including upregulation of sex hormone and WNT pathways and a downregulation of inflammation and cell cycle pathways. *Tnc^−^* mice had similar biomechanical properties after repair as WT. Further research is required to evaluate tissue specific alterations of *Tnc*, the interactions of *Tnc* and sex hormone and inflammation pathways as well as possible adjuvants to improve enthesis healing in the setting of reduced TNC function.

## Introduction

Failure of healing after rotator cuff repair (RCR) has been noted to occur on average between 15% to 20% of patients after surgical repair of the rotator cuff, although has been reported as high as 95%. (Longo 2021, Galatz 2004). A variety of genetic variants have been identified with rotator cuff tearing over the past 10 years, although very few have been clinically associated with impaired healing. (Tashjian 2021, Kluger 2017, Ball 2004, Tashjian 2016). Variants in both *Estrogen related receptor beta (ESRRB*) and *Tenascin-C* (*TNC*) have both been associated with inferior healing after RCR. (Kluger 2017, Tashjian 2016). None of the previously identified genes that have variants associated with either rotator cuff tears or impaired healing have been investigated *in vivo*.

TNC is a multi-modular, extracellular matrix protein with numerous molecular forms and functions with tightly controlled expression. (Midwood 2016) In adult humans, very little TNC is expressed in normal tissues but it is highly expressed in developing embryos as well as during wound healing or pathologic conditions including chronic inflammation and cancer. (Midwood 2016) TNC has a primary structural role through binding with fibronectin, collagen, and various proteoglycans. (Orend 2014) TNC also has cell-signaling roles impacting cell proliferation through activation of EGFR as well as inflammatory pathways through activation of TLR-4. (Swindle 2001, Midwood 2009) Finally, it can bind with soluble growth factors including FGF, PDGF, VEGF and TGF-beta which may affect their ability to signal to the cell. (Midwood 2016) Utilizing a murine RCR model we have determined that TNC is not expressed in native supraspinatus rotator cuff tendon, but it is induced around the tendon insertion site after repair, and sex hormone supplementation leads to a further upregulation of TNC. (Tashjian 2021, Tashjian 2024) Given the association of *TNC* variants with rotator cuff tearing and failure of healing and that TNC is upregulated at the repair site, we hypothesize that TNC may have a critical role in achieving healing after rotator cuff repair.

The purpose of the current study was to evaluate the effects of TNC on rotator cuff repair healing using our previously described murine rotator cuff repair model. (Tashjian 2021, Miller 2021, Tashjian 2024) Histologic, gene expression, animal activity and biomechanical evaluations were performed comparing Wild-Type, *Tnc* null, and *Tnc* heterozygous mice. We hypothesized that deletion or reduction of TNC would impair the histology, gene expression profiles, animal activity, and repair biomechanics.

## Materials and Methods

This study was approved by the University of Utah’s Institutional Animal Care and Use Committee (IACUC; Protocol number 21-07004).

### Animals and Surgery

Sixty male mice (12–14-week-old for RCR and 20-22-week-old for age-matched unrepaired controls) were utilized for all experiments. Overall, there were 21 wild-type animals (C57BL/6J; WT), 30 *Tenascin C* null mice (*Tnc^tm1Sia^; Tnc^−^*), and 9 *Tenascin C* heterozygous mice (*Tnc*^+/−^). (Saga 1992) Histologic, activity testing, RNA sequencing analysis and biomechanical tests were performed on operated (WT-RCR and *Tnc^−^-*RCR) mice, and histologic analysis was performed on *Tnc*^+/−^ mice. The unoperated, contralateral shoulders of the WT-RCR, *Tnc^−^-*RCR, and *Tnc*^+/−^ mice were used for histologic controls. Another group of age-matched unoperated WT and *Tnc^−^* mice were used for biomechanical controls. We used an established murine RCR model that was previously used for histological, cellular, molecular, and biomechanical analyses. (Tashjian JOR 2021, Tashjian JOR 2024, Miller 2021)

### Histologic Analysis

WT-RCR (n=3), *Tnc^−^-*RCR (n=7), and *Tnc^+/−^-*RCR (n=9) were sacrificed at 8 weeks postoperative and used for histologic analysis. The contralateral shoulders were utilized as controls. Mice were euthanized using isoflurane sedation followed by cervical dislocation. Shoulders were dissected and fixed in 10% neutral buffered formalin at 4°C for 48h and processed for histological analysis as previously described. (Tashjian JOR 2024) Slides were stained with toluidine blue or hematoxylin and eosin. Changes in histological parameters of the shoulder allowed us to quantify the degree of healing following RCR. We utilized a semi-quantitative scoring system described in Su *et al*. (Su AJSM 2018) Briefly, the following parameters were scored on a 1–4 scale and used to quantify histological changes: cellularity, proportion of cells resembling tenocytes, proportion of cells/fibers oriented parallel, and remodeling of the tendon-bone interface (scored by continuity, presence of fibrocartilage, regularity, and presence of a tidemark). Scoring was conducted by two blinded investigators and comparisons were made between the repaired and the contralateral unrepaired WT, *Tnc^−^*, and *Tnc^+/−^* shoulders.

### Animal Activity

WT-RCR (n=4) and *Tnc^−^-*RCR (n=8) underwent activity testing at 8 weeks postoperative. We used open field locomotor boxes with running wheels (SuperFlex Open Field Boxes; Omnitech Electronics, Inc.) to quantify changes in behavior/activity in mice. (Deacon 2006) We analyzed voluntary wheel running in control and experimental groups (Tashjian JOR 2024). The control and experimental groups were acclimated at 7 weeks post-RCR surgery to the open field locomotor boxes for 1 h on 4 consecutive days. Following 2 days’ rest, they were retested for 1 h on 4 consecutive days. Animals were handled by the same investigator and tested at the same time each day. Following completion of the behavioral/ activity analysis all animals were euthanized.

### Biomechanical Testing

WT-RCR (n=7), *Tnc^−^-*RCR (n=7), and unrepaired controls (4 WT and 4 *Tnc^−^*) were sacrificed and underwent biomechanical testing at 4 weeks postoperative evaluating stiffness and maximum loads. Biomechanical testing was performed on unrepaired controls and repaired WT- and *Tnc^−^-* mice. Animals were euthanized at 4 weeks post-RCR and the supraspinatus tendon was dissected under variable magnification stereo microscopy to isolate the supraspinatus tendon-bone construct. Tendon-bone constructs were tested for monotonic failure properties. (Lebaschi 2018) The muscle insertion was secured between pieces of waterproof sandpaper using cyanoacrylate glue. The humerus was gripped using a set of custom three-dimensional printed clamps with a cavity geometry specific to the mouse humerus and that positioned the humerus at a 60° abduction angle. (Kurtaliaj 2019) Hydration of the tendon and insertion was maintained by application of saline (0.9% NaCl) during dissection and test specimen preparation. Following application of a 0.1 N preload, samples were loaded to failure at 1 mm/min using a material test system (Electroforce 3300 Series II; TA Instruments). From these tests, maximal tangent stiffness and maximal load to failure were determined from the load-displacement data.

### RNA sequencing and enrichment analysis

WT-RCR (n=3) and *Tnc^−^-*RCR (n=4) were sacrificed at 2 weeks postoperative and used for RNA-seq. RCR was performed on seven mice (WT-RCR (n=3) and *Tnc^−^-*RCR (n=4)) and mice were euthanized 2 weeks post-RCR. Whole shoulder joints were exposed by removing the skin and the majority of muscle surrounding the joint with minimal disruption of the ligaments, tendons, and cartilage. The humerus and scapula were cut approximately 1 cm distal from the articular surface. Joints were transferred to new RNAlater solution, rinsed, cut into small fragments, transferred to a tube containing Trizol and 2.8mm stainless steel beads, and homogenized using the BeadBug microtube homogenizer. Total RNA was isolated using a Direct-zol RNA Miniprep kit (Zymo). RNA-seq, quality control, and alignment was performed by Novogene. Aligned reads were counted using HTSeq v0.11.3, and the counts were then analyzed for differential gene expression using median-ratio-normalization with Deseq2 v1.30.0 as previously described. (Jurynec 2022) Fold-changes were calculated by comparing read counts in *Tnc^−^-*RCR joints relative to WT-RCR joints. Genes with an adjusted P-value < 0.05 were considered differentially expressed. We used Ingenuity Pathway Analysis (Qiagen) software for functional pathway analysis of the differentially expressed genes identified in this study. The log2 fold change and adjusted p values using Bonferroni method from Deseq2 analysis were used to calculate directionality (Z score) and set canonical pathway p-values (<0.05). Data has been deposited to the Gene Expression Omnibus (GSE271514).

### Statistical analysis

#### Histological Analysis

Statistical significance was determined by two-way ANOVA with Tukey’s multiple comparisons test (Figure 1) and by a two-tailed unpaired t-test (Figure 2). Error bars in Figures 1 and 2 represent mean ± SD and statistically significant differences of **p ≤ 0.01.

**Figure 1.**
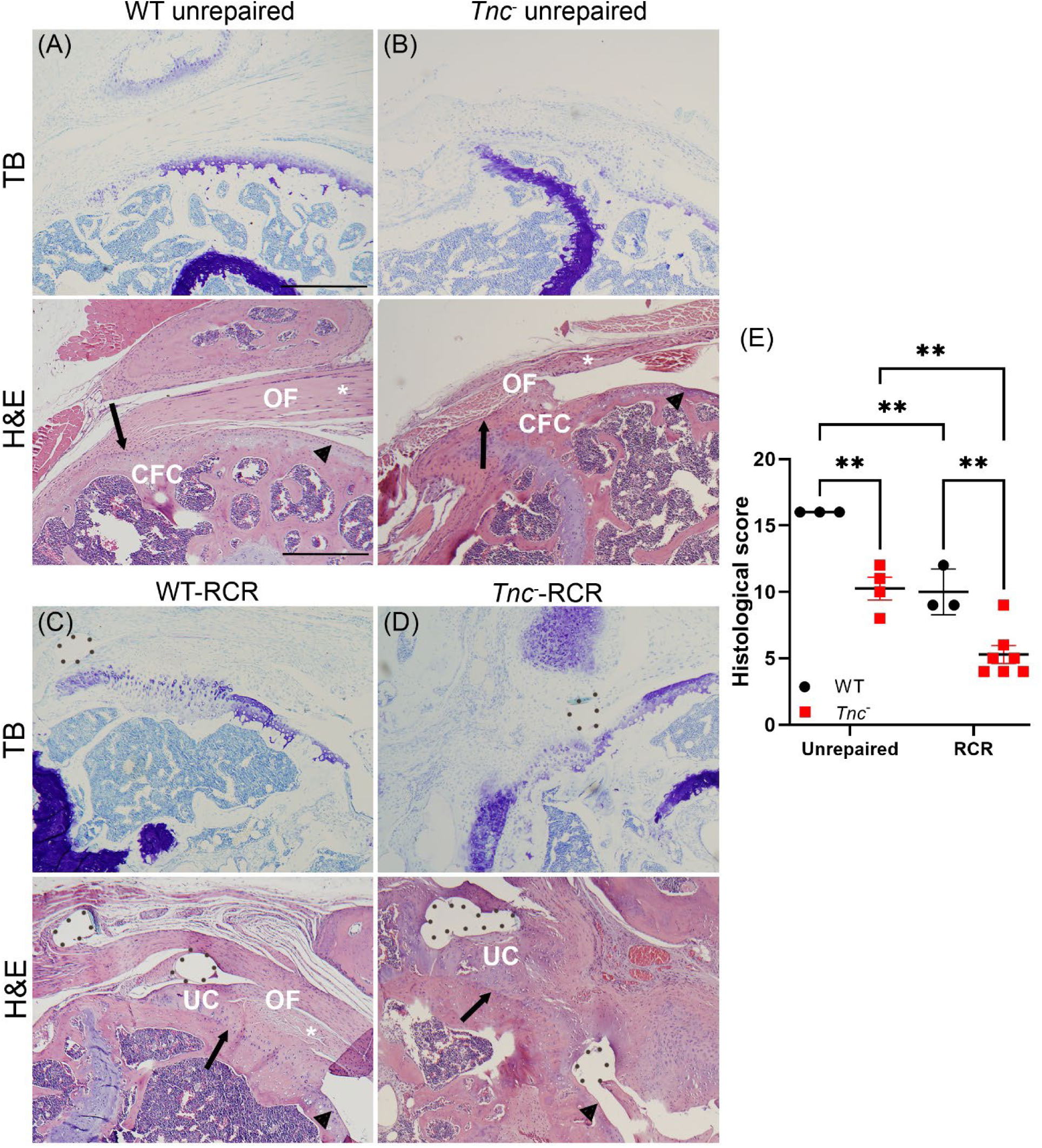
*Tenascin C* null mice have severe bone and tendon defects following rotator cuff repair. Histological analysis of shoulder tissue sections 8 weeks post-RCR. Coronal sections of the shoulder joint with the humerus to the left and scapula to the right. (A and B). Unrepaired WT and *Tnc^−^ s*houlders indicating supraspinatus tendon attachment to the humeral head (arrow). Toluidine blue (TB) staining marks the supraspinatus enthesis and hematoxylin and eosin staining (H&E) indicates the structure of the organizing fibers (OF), calcified fibrocartilage (CFC), and the supraspinatus tendon (asterisks). Histologial structure of the WT unrepaired (A) and *Tnc^−^*unrepaired shoulders (B). (C) WT-RCR shoulders have increased CFC and disorganization of the OF compared with WT unrepaired shoulders (A). (D) *Tnc^−^-*RCR animals have a high degree of CFC, severe disorganization of the organizing fibers, increased calcified cartilage (UC) and increased bone remodeling compared with unrepaired controls (A and B) and WT-RCR animals (C). Dashed lines indicate suture insertion through the supraspinatus tendon. Arrows mark the tendon-bone interface and asterisks mark the supraspinatus tendon, which were used to quantify histological changes (cellularity, proportion of cells resembling tenocytes, and proportion of cells/fibers oriented parallel in the supraspinatus tendon and remodeling of the tendon-bone interface). Arrowheads mark the articular surface of the humeral head. (E) Graph indicates quantification of histological data. WT unrepaired; n = 3, *Tnc^−^*unrepaired; n = 3, WT-RCR; n = 3, *Tnc^−^*-RCR; n = 7. n = independent biological replicates. Error bars represent mean ± SD and statistically significant differences of **p ≤ 0.01. Scale bars = 250µm.

**Figure 2.**
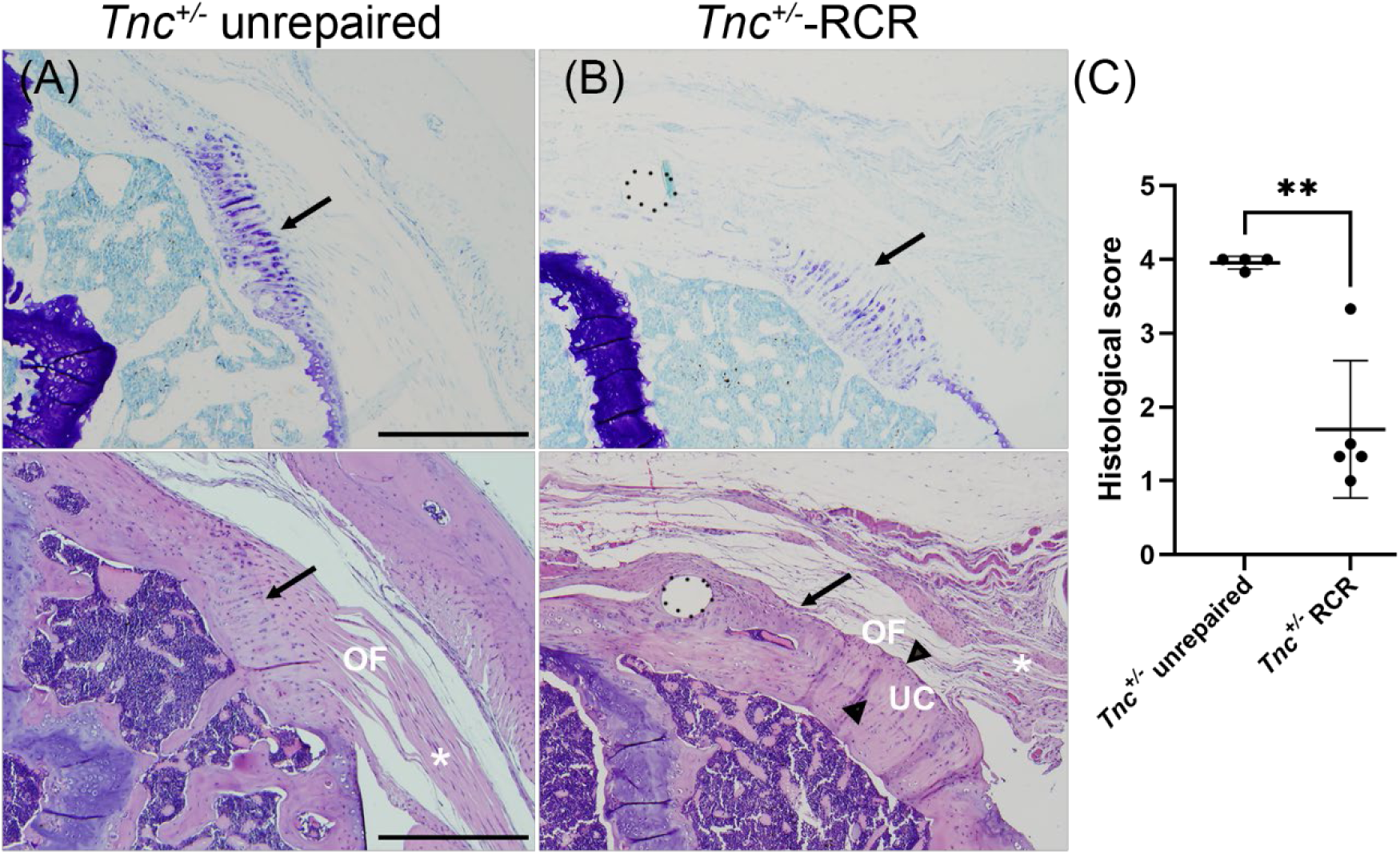
*Tenascin C* heterozygous mice have tendon defects following rotator cuff repair. Histological analysis of shoulder tissue sections 8 weeks post-RCR. Coronal sections of the shoulder joint with the humerus to the left and scapula to the right. (A and B). Unrepaired *Tnc^+/−^* (A) and *Tnc^+/−^* -RCR (B). Toluidine blue (TB) staining marks the supraspinatus tendon and enthesis and hematoxylin and eosin staining (H&E) indicates the structure of the organizing fibers (OF), calcified cartilage (UC), and the supraspinatus tendon (asterisks). (A) Unrepaired *Tnc^+/−^* shoulders (the contralateral shoulder from RCR mice) are structurally indistinguishable from WT unprepared shoulders. (B) *Tnc^+/−^* -RCR shoulders have a severe disorganization of the organizing fibers and supraspinatus tendon and increased formation of uncalcified cartilage (UC, marked by arrowheads) compared with unrepaired *Tnc^+/−^* controls. Dashed lines indicate suture insertion through the supraspinatus tendon. Arrows mark the tendon-bone interface and asterisks mark the supraspinatus tendon, which were used to quantify histological changes (cellularity, proportion of cells resembling tenocytes, and proportion of cells/fibers oriented parallel in the supraspinatus tendon and remodeling of the tendon-bone interface). (C) Graph indicates quantification of histological data. Unrepaired *Tnc^+/−^*; n = 4, *Tnc^+/−^* -RCR; n = 5. n = independent biological replicates. Error bars represent mean ± SD and statistically significant differences of **p ≤ 0.01. Scale bars = 250µm.

#### Activity Testing

We used the Shapiro–Wilk test to assess normality of the data and Grubb’s test with α = 0.05 to detect and remove outliers. Statistical significance was determined by a two-tailed unpaired t-test. Error bars in Figure 3 represent mean ± SD and statistically significant differences of *p≤ 0.05, **p ≤ 0.01 and ****p ≤ 0.0001.

**Figure 3.**
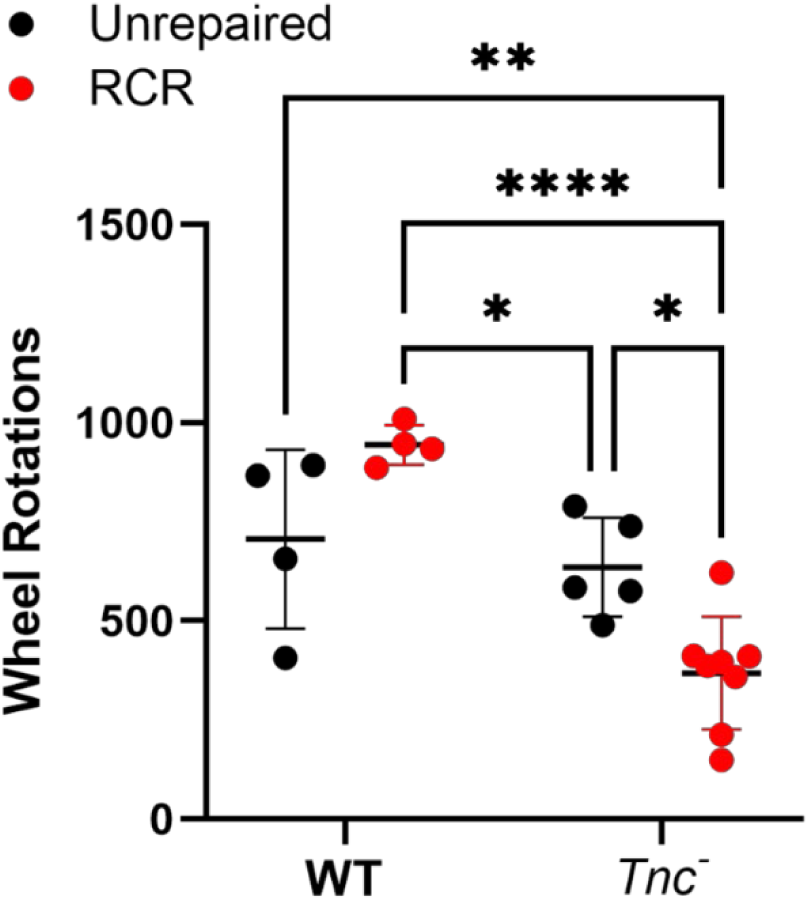
*Tenascin C* null mice have reduced wheel rotations after rotator cuff repair. Graphs represent 4-day averages for each animal. Graphs represent the number of wheel rotations. Unrepaired WT; n =4, unrepaired *Tnc^−^*; n =5, WT-RCR; n = 4 and *Tnc^−^*-RCR; n = 8. N = independent biological replicates. Error bars represent mean ± SD and statistically significant differences of *p≤ 0.05, **p ≤ 0.01 and ****p ≤ 0.0001.

#### Biomechanical Testing

A power and sample size analysis using maximum load estimates from pilot data indicated that 11 samples per group would provide 80% power to test equivalence in a ±1.0N range. Grubb’s test with α = 0.05 was used to detect and remove outliers for both maximum load and stiffness. Two-tailed unpaired t-tests were used to test for differences between unrepaired controls and repair groups, and between RCR groups, with Holm-Sidak correction for multiple comparisons. A significance level of 0.05 was used for all tests.

## Results

Given the association of *TNC* variants with inferior healing after RCR (Kluger 2017, Tashjian 2016) and the positive regulation of *Tnc* by sex hormones in shoulder tissue after RCR (Tashjian JOR 2024), we wanted to test if loss of TNC affected healing after RCR. We first analyzed the histological structure of unrepaired WT and *Tenascin C* null (*Tnc^−^*) shoulders. (Figure 1A, B, and E) Qualitative histological analysis of unrepaired 20–23-week-old shoulders indicated that *Tnc^−^* mice had mild structural defects (reduced tendon size and increase in calcified fibrocartilage) in the supraspinatus tendon and enthesis compared with WT. (Figure 1A and B) Semi-quantitative assessment of the histology indicated a reduction in overall histological score. (Figure 1E). We next performed RCR on WT and *Tnc^−^* mice. Qualitative histological analysis at 8 weeks post-RCR demonstrated that *Tnc^−^* mice had severe bone and tendon defects following rotator cuff repair (RCR). *Tnc^−^-*RCR animals had a high degree of uncalcified fibrocartilage, severe disorganization of the organizing fibers, and increased bone remodeling compared to unrepaired contralateral WT and *Tnc^−^* shoulders (Figure 1A-D). Semi-quantitative assessment of the histology supported significantly greater disorganization of the tendon enthesis of unrepaired *Tnc^−^* animals compared to WT animals as well as significantly greater disorganization of *Tnc^−^-* RCR animals compared to both WT-RCR and unrepaired *Tnc^−^* control animals. (Figure 1E)

*TNC* variants have a dominant effect on inferior healing after RCR. (Kluger 2017, Tashjian 2016) Given that unrepaired *Tnc^−^* shoulders had mild structural defects (Figure 1A, B, and E) we wanted to determine if shoulders from unrepaired heterozygous *Tnc* mice (*Tnc^+/−^*) were structurally normal and if loss of one copy of *Tnc* was sufficient to alter the cellular response to RCR. We performed RCR on *Tnc^+/−^* mice and analyzed the histological structure of the repaired and contralateral shoulders 8 weeks post-RCR. Qualitative histological analysis and semi-quantitative assessment of the unrepaired contralateral shoulder indicated that *Tnc^+/−^* were structurally indistinguishable from unrepaired WT controls. (Compare Figure 1A and E with Figure 2A and C). In response to RCR, shoulders from *Tnc^+/−^* -RCR mice displayed a severe disorganization of the organizing fibers and supraspinatus tendon and had an increased formation of uncalcified cartilage compared with unrepaired *Tnc^+/−^*contralateral shoulders. (Figure 2A-C). In sum, these data indicate that TNC is needed to mount a normal repair response to RCR and that loss of one copy of *Tnc* is sufficient to alter the normal cellular response to RCR.

Next, we wanted to determine if the histological changes observed in *Tnc^−^* mice are associated with changes in animal behavior. (Tashjian JOR 2024) To test this, we analyzed wheel activity using unrepaired WT, unrepaired *Tnc^−^*, WT-RCR, and *Tnc^−^*-RCR mice. There no differences in wheel rotations between the unrepaired WT and *Tnc^−^* mice. In contrast, *Tnc^−^*-RCR mice had a reduced number of wheel after rotator cuff repair with WT-RCR (Figure 3). These data indicated the severe bone and tendon remodeling in *Tnc^−^*-RCR mice correlated with a reduction in activity.

Given the structural and behavioral defects associated with loss of TNC function, we wanted to test if these correlated with changes in tendon biomechanics. At 4-weeks post-RCR, we examined the max load and max stiffness in unrepaired WT, unrepaired *Tnc^−^*, WT-RCR, and *Tnc^−^*-RCR tendons. RCR in both the *Tnc^−^* and WT mice resulted in a significant reduction in overall biomechanical properties with significant reductions in both maximal load to failure (WT-unrepaired vs. WT-RCR, p=0.014; *Tnc^−^*unrepaired vs. *Tnc^−^-*RCR, p<0.007) and stiffness (WT-unrepaired vs. WT-RCR, <0.001; *Tnc^−^* unrepaired vs. *Tnc^−^*-RCR, p<0.001) Loss of *Tnc* did not have a significant effect on repair biomechanics comparing *Tnc*^−^-RCR and WT-RCR (all p>0.6) (Figure 4A and B).

**Figure 4.**
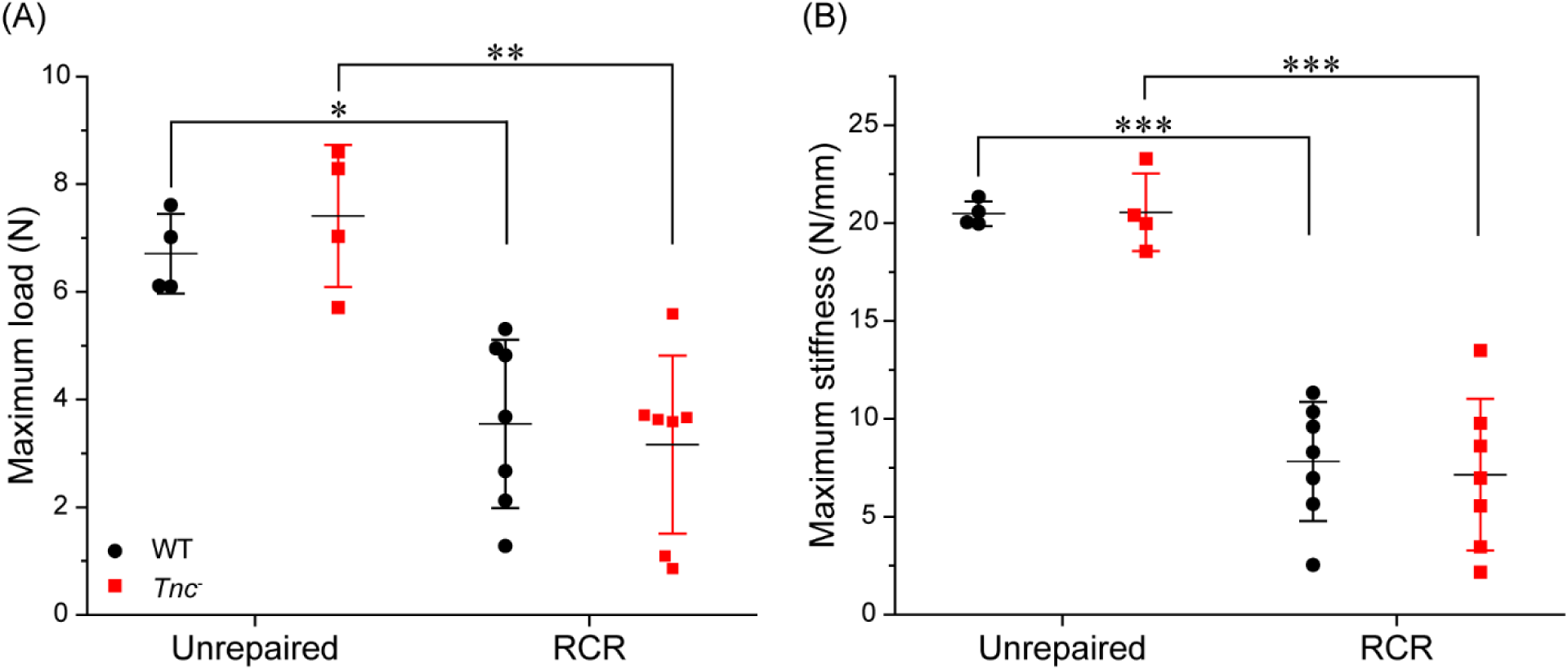
Tendons isolated from *Tenascin C* null mice do not have altered biomechanical properties. (A) *Tnc^−^* tendons had similar ultimate failure loads to WT mice after repair (Maximum load (N)). There was no difference within unrepaired control or repaired groups. (B) *Tnc^−^* tendons had similar stiffness to WT mice after repair (Maximum stiffness (N/mm)). There was no difference within unrepaired control or repaired groups (p=0.923). Dots are independent biological replicates. Error bars represent mean ± SEM and statistically significant differences of *p ≤0.05, **p ≤ 0.01, and ***p ≤0.001.

Finally, we sought to identify the early molecular changes in the shoulder that occur after RCR in WT and *Tnc^−^* mice. Loss of TNC following RCR significantly altered gene expression in the shoulder compared with WT-RCR. (Figure 5) At 14 days post-RCR, loss of *Tnc* led to upregulation of the estrogen and androgen receptors (*Esr1, Esr2,* and *Ar*). (Figure 5A) Similarly, many WNT pathways were upregulated in the *Tnc^−^*-RCR shoulders, including *Wnt2, Wnt9a, Wnt11* and *Fzd4*. Furthermore, we noted an upregulation of *Wnk2*, which is a protein involved in regulating the response to hyperosmotic stress. (Figure 5A). To discover pathways regulated by TNC function we used IPA software to identify the top-ranking pathways determined by Z-score, which predicts the significance and directionality of pathway activation or inhibition. Top pathways upregulated in *Tnc^−^*-RCR shoulders are those involved in mitochondrial function and metabolism, estrogen and testosterone signaling, and calcium signaling. (Figure 5B) The top pathways downregulated by loss of TNC function include several proinflammatory pathways (e.g., IL-17, IL-2, and IL-3, IL-5 and GM-CSF signaling) and those associated with a reduction in the cell cycle (e.g., synthesis of DNA, and cell cycle checkpoints). (Figure 5B)

**Figure 5.**
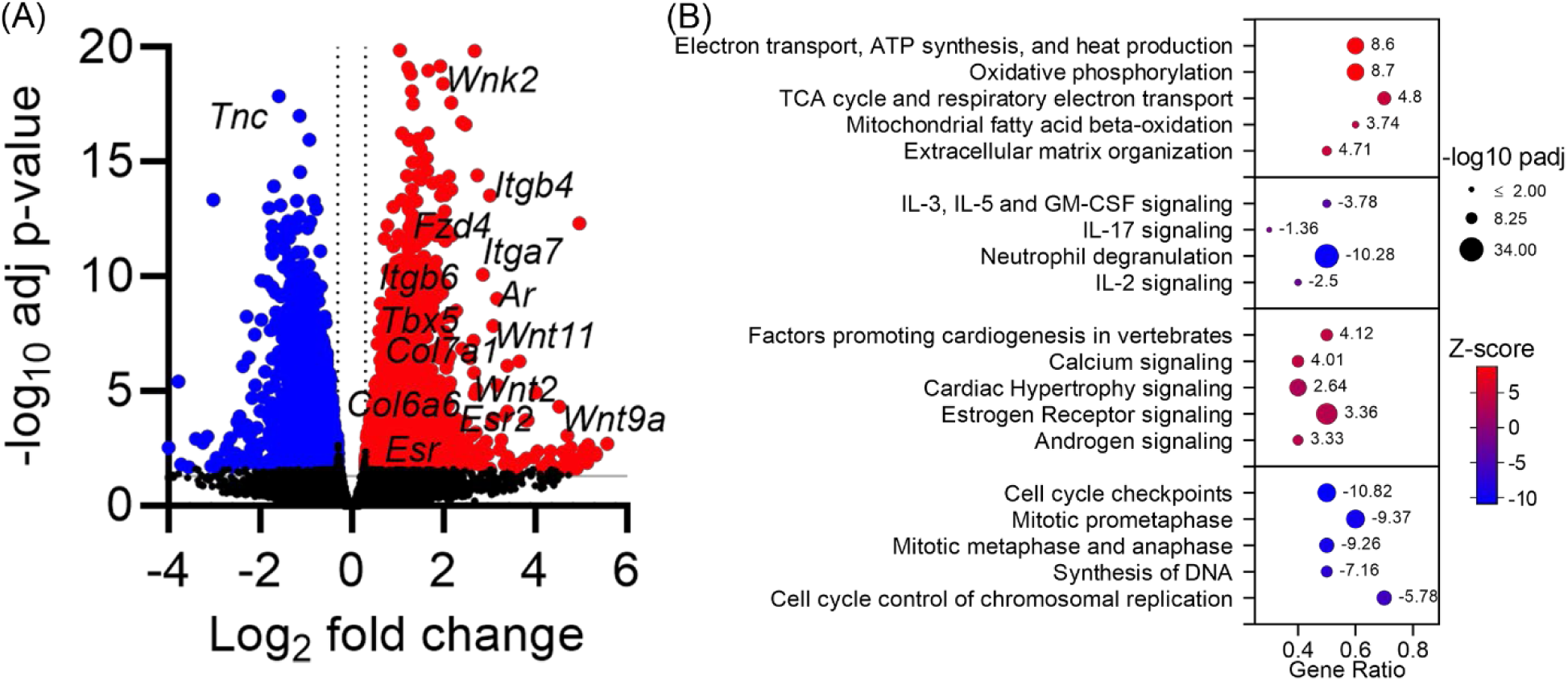
Loss of *Tenascin C* following rotator cuff repair altered gene expression in the shoulder. RNA-seq analysis of RNA isolated from WT and *Tnc^−^* shoulders 14 days post-RCR. (A) Volcano plot indicates genes significantly upregulated (red) or downregulated (blue) in *Tnc^−^*shoulders compared with WT-RCR shoulders. (B) Z-scores indicate pathways that are up- or downregulated in *Tnc^−^-*RCR compared with WT-RCR shoulders. WT-RCR; n = 3 and *Tnc^−^*-RCR; n = 4 for each genotype. n = independent biological replicates.

In conclusion, our data indicate the TNC is necessary for the normal cellular and molecular repair response to RCR. These underlying changes are correlated with a reduction in animal activity, but not with changes in tendon biomechanics.

## Discussion

Deletion of *Tenascin C (Tnc)* results in a disorganized enthesis after rotator cuff repair, reduces activity, and alters gene expression when compared to repairs in control animals although no differences in the biomechanical strength of rotator cuff repairs resulted from *Tnc* deletion. Histologically, *Tnc* deletion leads to increased uncalcified cartilage and fiber disorganization, even in comparison to control repairs where uncalcified cartilage is increased in comparison to the normal tendon enthesis. Animal activity was dramatically reduced in *Tnc^−^*-RCR compared to control RCR supporting the hypothesis that that impaired tendon enthesis restoration results in reduced animal function. Upregulation of estrogen receptor and androgen receptor signaling pathways support a failure of hormone functionality on tendon healing and downregulation of cell cycle and inflammatory pathways support dysregulation of key phases (inflammation, reparative) of tendon healing. Overall, the data supports the hypothesis that *TNC* variants associated with impaired healing likely led to a reduction in TNC function and may have a direct causative effect on rotator cuff repair failure.

Several biological pathways are altered in *Tnc^−^-*RCR mice compared with WT-RCR mice including sex hormone receptors (Tashjian JOR 2024, Tashjian JOR 2021), 1L-17 signaling, and neutrophil degranulation. Our RNA-seq data indicated that there is complex cellular interaction during the repair process. There is a well-established balance of pro- and anti-inflammatory processes that are needed during regenerative and healing processes in many tissues, including tendon. IL-17 is a key proinflammatory mediator. It promotes tissue destruction during inflammation and it induces the production of IL-1, TNF-alpha, and matrix metalloproteinases by fibroblasts and macrophages. (Millar 2016) Reduction of IL-17 production in the setting of tendon healing may halt the process of repair by reducing the early proinflammatory response to injury. Upregulation of energy metabolism pathways in the setting of downregulation of cell cycle pathways would support increased efforts to repair or rebuild but an inability to do so secondary to the impaired ability to replicate cells. As a result, normal tendon repair and remodeling which occurs after inflammation is not allowed to occur. Typically, monocytes are transformed to support new tissue formation but without the ability to replicate this process would be impaired. (Zumstein 2017)

Estrogen receptor and androgen receptor pathways were upregulated in *Tnc^−^*-RCR shoulders. This would support that appropriate hormonal signaling is not occurring in the setting of TNC deficiency. We have previously shown that estrogen supplementation after rotator cuff repair promotes growth and repair through PDGF and TGF-beta pathways and reducing inflammation while testosterone positively regulates genes associated with muscle contraction and myoblast fusion. (Tashjian JOR 2024) Improved activity levels were also noted in animals supplemented with hormones, specifically testosterone, therefore the impaired activity noted in the TNC deficient animals may be resultant of impaired hormone pathways. (Tashjian JOR 2024) Further research is required to evaluate tissue specific alterations of TNC activity (tendon, bone, etc) and the interaction of TNC and sex hormone signaling. More data is needed to determine if we can develop adjuvants to improve enthesis healing in the setting of reduced TNC activity in patients undergoing RCR.

*Tenascin C* deletion resulted in upregulation of several WNT ligands and receptors after repair. The WNT pathway plays a critical role in pathological calcification and may result in heterotopic ossification in tendon tissue. (Liu 2015) WNT/β-catenin signaling also suppresses scleraxis and tenomodulin which are critical for tendon maturation and eventual healing. (Kishimoto 2015, Gulotta 2011,Tokunaga 2015) Increased heterotopic ossification in the tendon tissue along with downregulation of normally expressed tendon modulating genes may be one potential cause for the abnormal histologic appearance with areas of increased bone remodeling. Restoration of a normal tendon enthesis is the goal of tendon repair surgery and therefore impairment of TNC function can result in grossly abnormal anatomy with increased ossification.

Lack of differences in biomechanical analysis may be a result from failure to define the optimal timing for analysis. Further time points would need to be evaluated to determine if timing of sacrifice influences mechanical testing. Most importantly, because rotator cuff repairs heal so quickly and reliably in this small animal model, the clinical translatability of the results of mechanical testing to the human condition are unclear. Large animal studies are required to determine if *Tnc* deletion results in significantly impaired biomechanics that would be clinically important.

Our study has several limitations. We used a global knockout of *Tnc* to examine the role of TNC during the healing process following RCR. Without tissue specific deletion of *Tnc*, we are unable to determine which tissues in the shoulder require TNC function for proper healing following RCR. Furthermore, we are unable to determine precisely which tissues in the shoulder have altered gene expression following RCR. The changes may be in local tissues (tendon or bone) or other cell types such as invading immunomodulatory or systemic factors, especially since we observed a downregulation of several inflammatory pathways in the shoulder of *Tnc^−^* mice after RCR. Regardless of where TNC is required, our data clearly demonstrate the TNC function is necessary for healing following RCR. This is consistent with the association of *TNC* variants with inferior healing after RCR in humans. (Kluger 2017, Tashjian 2016) The generation of a conditional allele of *Tnc* will allow us to determine the spatial and temporal requirements of TNC during the repair process. We only utilized male mice in this study. We do not know if healing following RCR is different in males vs females. Sex is an important biological factor to consider, especially in the context of therapeutic treatments. We note that our previous study demonstrated the systemic supplementation of male mice with estrogen or testosterone following RCR significantly improved healing (Tashjian 2023), indicating that sex specific hormones may be a potential therapeutic option to improve patient outcomes following RCR.

In conclusion, our data supports that *Tnc* deletion significantly impairs rotator cuff repair histologic properties, animal function and alters gene expression in comparison with WT controls. Future studies are required to determine the precise tissue and cellular requirement of TNC and if biologic augmentation may improve healing in the setting of reduced TNC function.

## Notes

**Sources of Support:** Funding for this project was provided by the LS Peery Foundation Discovery Program in Musculoskeletal Restoration, the Skaggs Foundation for Research (MJJ), the Arthritis National Research Foundation (MJJ – 707634), the National Institute on Aging (MJJ – R21AG063534), and National Institute of Arthritis and Musculoskeletal and Skin Disease (MJJ and NHK - R01AR082973).

